# Host-induced silencing of the *Colletotrichum gloeosporioides conidial morphology 1* gene (*CgCOM1*) confers resistance against Anthracnose disease in chilli and tomato

**DOI:** 10.1101/2020.01.03.893677

**Authors:** Binod Kumar Mahto, Anjulata Singh, Manish Pareek, Manchikatla V. Rajam, Swatismita Dhar-Ray, Pallavolu M. Reddy

## Abstract

Anthracnose disease is caused by the ascomycetes fungal species *Colletotrichum,* which is responsible for heavy yield losses in chilli and tomato worldwide. Conventionally, harmful pesticides are used to contain anthracnose disease with limited success. In this study, we assessed the potential of Host-Induced Gene Silencing (HIGS) approach to target the *<u>C</u>olletotrichum gloeosporioides COM1* (*CgCOM1*) developmental gene involved in the fungal conidial and appressorium formation, to restrict fungal infection in chilli and tomato fruits. For this study, we have developed stable transgenic lines of chilli and tomato expressing *CgCOM1*-RNAi construct employing *Agrobacterium*-mediated transformation. Transgenic plants were characterized by molecular and gene expression analyses. Production of specific *CgCOM1* siRNA in transgenic chilli and tomato RNAi lines was confirmed by stem-loop RT-PCR. Fungal challenge assays on leaves and fruits showed that the transgenic lines were resistant to anthracnose disease-causing *C. gloeosporioides* in comparison to wild type and empty-vector control plants. RT-qPCR analyses in transgenic lines revealed barely any *CgCOM1* transcripts in the C. *gloeosporioides* infected tissues, indicating near complete silencing of *CgCOM1* gene expression in the pathogen. Microscopic examination of the *Cg*-challenged leaves of chilli-*CgCOM1*i lines revealed highly suppressed conidial germination, germ tube development, appressoria formation and mycelial growth of *C. gloeosporioides*, resulting in reduced infection of plant tissues. These results demonstrated highly efficient use of HIGS in silencing the expression of essential fungal developmental genes to inhibit the growth of pathogenic fungi, thus providing a highly precise approach to arrest the spread of disease.

## Introduction

Chilli and tomato are widely grown Solanacious crops, both in tropical and subtropical regions of the world. Anthracnose is a major disease of chilli and tomato. It causes pre- and post-harvest necrotic lesions mainly on fruits leading to yield losses up to 50% (Bosland and Votava, 2012; De Silva *et al.,* 2017; Hyde *et al.,* 2009; Kim *et al.,* 2008; Pakdeevaraporn *et al.,* 2005; Phoulivong *et al.,* 2010; Prathibha *et al.,* 2013; Than *et al.,* 2008). Anthracnose disease is caused by ascomycete fungi belonging to the genus *Colletotrichum,* which consists of approximately 600 species including most studied species like *C. gloeosporioides, C. capsici, C. acutatum, C. dematium, C. nigrum* and *C. coccodes*. *Colletotrichum* species is known to infect more than 3200 species of monocot and dicot plants (O’Connell *et al.*, 2012). From among these fungal species, *C. gloeosporioides* is able to infect and cause disease in both chilli and tomato (Barksdale, 1972; Li, 2013; Than *et al.*, 2008). *C. gloeosporioides* is a hemibiotrophic fungus, which exhibits a short biotrophic phase, followed by a longer necrotrophic phase on the host plant (Alkan *et al.,* 2015; O’Connell *et al.,* 2012). During the biotrophic phase, fungal species develop specialized infection structures such as germ tubes and melanized appressoria on host surface. Appressoria produce thin primary penetration hyphae (intracellular hyphae), which infiltrate the host plasma membrane and give rise to secondary necrotrophic hyphae that cause visible disease symptoms in plants (Barksdale, 1972; Jeger and Bailey, 1992; Nelson, 2008; O’Connell *et al.,* 2004; O’Connell *et al.,* 2012; Takahara *et al.,* 2009; Than *et al.,* 2008; Voorrips *et al.,* 2004).

Currently, integrated pest/disease management techniques comprising of chemical, physical and biological strategies are being employed to control anthracnose disease in chilli and tomato crops, but with limited success (Than *et al.*, 2008). Even conventional breeding techniques have not been successful in developing effective resistant varieties against this fungal disease (Than *et al.*, 2008). So, this has prompted scientists to adopt new technologies such as transgenic approach to develop resistant plant varieties. For example, Seo *et al.* (2014) overexpressed the plant defensin gene *J1-1* to neutralize the pathogen and control the spread anthracnose disease in fruits of chilli pepper. However, this approach conferred only 50% resistance against this disease.

Lately, RNAi technique is being used to target plant pathogens to control the spread of many diseases in plants (Yogindran and Rajam, 2015). RNAi is a post transcriptional gene silencing (PTGS) mechanism where siRNA, generated through the actions of dicer/RISC, mediates degradation of target mRNA (Baulcombe, 2004; Vaucheret *et al.,* 2001). Host Induced Gene Silencing (HIGS) is a novel application of RNAi in which the plants are genetically modified to express siRNA to target pathogen genes that are crucial for the development, pathogenicity and survival of the pest on plants. Earlier studies have shown that the HIGS can be used as a useful tool to defend plants from nematode infection (Dubreuil *et al.*, 2007; Huang *et al.*, 2006), insect attack (Baum *et al.*, 2007; Mao *et al.*, 2007; Reddy and Rajam, 2015) and fungal pathogens (Chen *et al.*, 2016; Cheng *et al.*, 2015; Ghag *et al.*, 2014; Govindarajulu *et al.*, 2015; Jahan *et al.*, 2015; Koch *et al.*, 2013; Panwar *et al.*, 2017; Panwar *et al.*, 2013; Song and Thomma, 2018; Thakare *et al.*, 2017; Zhou *et al.*, 2016). Hence, in the current study, we employed HIGS technology to control infection and spread of anthracnose disease-causing *C. gloeosporioides* in chilli and tomato, to render these crop plants free from the pathogen attack.

Recently, a Conidial Morphology 1 protein (COM1), was shown to play a critical role in conidial development and appressorial penetration in *Magnaporthe oryzae*, an ascomycetes pathogen, which causes devastating blast disease in rice (Bhadauria *et al.,* 2010; Yang *et al.,* 2010). Since *C. gloeosporioides* is also an ascomycetes species, we conjectured that an orthologous COM1 protein may likewise function in a similar manner in controlling the developmental processes in this fungal pathogen as well. Thus, we envisaged that the trans-expression of *CgCOM1* RNAi construct in the host chilli and tomato lines will produce *CgCOM1*siRNA, which upon taken up by *C. gloeosporioides* during the infection process will target the fungal *CgCOM1* mRNA leading to its degradation; thus inhibiting the development appresorial/infection apparatus in this fungal pathogen, and resulting in the prevention of manifestation of anthracnose disease in the plant. In the present study, we demonstrated that the host-induced silencing of *COM1* gene of *C. gloeosporioides* effectively restricts infection and spread of anthracnose disease in chilli and tomato.

## Material and methods

### Fungus strain and plant material

Fungal strain of *Colletotrichum gloeosporioides* (ITCC No. 6270) was obtained from Indian Type Culture Collection (ITCC), IARI, New Delhi, and grown in potato dextrose agar (PDA) medium at 28 °C.

Certified seeds of Indian cultivated chilli (*Capsicum annuum* L.), varieties Pusa Jwala (PJ) and Pusa Sadabahar (PSB), and tomato (*Solanum lycopersicum*) varieties Pusa Rohini (PR) and Pusa Early Dwarf (PED) were obtained from National Seeds Corporation (NSC), IARI, New Delhi, India.

### Preparation of fungal inoculum and infection assay

Fungal strain of *C. gloeosporioides* was grown on Potato Dextrose Agar (PDA) plates at 28°C. Spores were harvested from 3-day old cultures and spore inoculum (1×10^5^ spores ml^-1^) was prepared in sterile distilled water (Khatri and Rajam, 2007).

### *Construction of* CgCOM1*-RNAi vector and transformation of chilli and tomato*

Sense and anti-sense *CgCOM1* gene fragments were PCR-amplified using the sense (forward: 5’-GGCGCGCCAAGAAGCGCAAGGCC-3’ and reverse: 5’-ATTTAAATATGTCGGCAATTTC-3’) and antisense (forward: 5’-TCTAGAAAGAAGCGCAAGGCC-3’ and reverse: 5’-GGATCCATGTCGGCAATTTC-3’) primers, gel purified, ligated into *Sma1* digested pBlueScript SK (Stratagene, USA) vector, and sequence verified for fidelity (Xceleris, Ahmedabad, India) and utilized for the development of RNAi vector construct.

The sequence-verified pBSK-*CgCOM1*-sense and pBSK-*CgCOM1*-antisense clones were digested with *Asc*1/*Swa*1 and *Xba*1/*BamH*1 restriction enzymes, respectively, and purified using gel extraction kit (Qiagen, USA). Subsequently, first, *Asc*1/*Swa*1 digested sense fragment was cloned downstream of 35S CaMV promoter in a similarly restricted pMVR-hp binary vector (Reddy and Rajam, 2016), resulting in pMVR-hp-*CgCOM1*-S. Thus developed pMVR-hp-*CgCOM1*-S vector was later utilized to clone anti-sense *CgCOM1* fragment into it. The *Xba*1/*BamH*1 digested anti-sense fragment was ligated to likewise restricted pMVR-hp-*CgCOM1*-S vector to develop pMVR-hp-*CgCOM1i* plant transformation vector. *CgCOM1*-RNAi construct was mobilized into *Agrobacterium tumefaciens* LBA4404 strain through freeze thaw method (Hofgen and Willmitzer, 1988) and utilized for chilli and tomato transformation. Chilli and tomato transformation were carried out according to the protocols describe by Mahto *et al.* (2018) and Madhulatha *et al.* (2007), respectively.

### Molecular characterization of chilli and tomato transformants

The total genomic DNA was isolated from leaf tissue of putative *CgCOM1*-RNAi chilli and tomato lines using CTAB method (Doyle and Doyle, 1990) and employed for molecular confirmation.

Transgenic plants transformed with pMVR-hp-*CgCOM1*i construct and empty pMVR-hp vector were confirmed by PCR using *CgCOM1* gene or *CaMV35S* promoter-specific primers (Supplementary Table S1).

Transgene expression in chilli and tomato transgenic lines was analysed by RT-PCR and RT-qPCR. For expression analysis total RNA was extracted from 100 mg of freeze-dried leaf tissue of chilli and tomato lines using RNeasy plant kit (Qiagen, USA) according to manufacturer’s protocol. Concentration of total RNA was quantified using Nanodrop2000 spectrophotometer (Thermo Scientific, USA). The total RNA was reverse transcribed using Revert Aid H Minus first strand cDNA synthesis kit (Thermo scientific, USA) following the instructions of the manufacturer. Reverse transcribed cDNA was utilized for *CgCOM1* expression analysis using *CgCOM1* gene specific primers.

Relative expression of *CgCOM1* transgene in chilli- and tomato-*CgCOM1*i transgenic lines was analyzed by RT-qPCR using *CgCOM1* gene specific primers (Supplementary Table S1) and SSoFast™ EvaGreen^®^ supermix (Bio-Rad, USA) on CFX96 Touch™ Real-Time PCR detection system (Bio-Rad, USA) according to manufacturer’s instructions. Expression of *CgCOM1* gene was normalized with *CaGAPDH* of and *SlACTIN* expression in *C. annuum* and *S. lycopersicum*, respectively. The relative changes in gene expression were computed by 2^−ΔΔCt^ method (Livak and Schmittgen, 2001).

### Detection of siRNA in transgenic chilli and tomato

To detect *CgCOM1* siRNA production in chilli- and tomato-*CgCOM1*i transgenic plants, stem-loop RT-PCR (Varkonyi-Gasic *et al.*, 2007) was performed against the *CgCOM1* gene of *C. gloeosporioides*. For this purpose, initially through *in silico* analysis (www.sirna.wi.mit.edu), putative siRNA sequences were identified within the 168 bp *CgCOM1* sequence employed for the construction of RNAi vector (Supplementary Fig. S1), and scored according to Reynolds *et al.* (2004) to pinpoint/identify the potential siRNA, which is functionally appropriate for knocking down the *CgCOM1* gene expression. The siRNA sequence that gave the highest score was selected for designing the siRNA specific forward primer. For achieving the ampification of 60 bp amplicon, first low molecular weight RNA was isolated from total leaf RNA according to Peng *et al.* (2014). Low molecular weight RNA was then utilized to synthesize cDNA using stem-loop primer, designed according to Chen *et al.* (2005) (Supplementary Table 1). Subsequently, stem-loop RT-PCR was performed using siRNA specific forward (see above) and universal reverse primers to amplify 60 bp amplicon encompassing siRNA from transgenic leaves. Low molecular weight RNA isolated from wild-type plant served as a control.

### *Fungal challenge assays on chilli- and tomato-*CgCOM1*i transgenic plants and* determination of CgCOM1 *transcript abundance in infected tissues*

*C. gloeosporioides* challenge assays were performed on detached 5-week old leaves and fruits of chilli-RNAi (Pusa Jwala-*CgCOM1*i and Pusa Sadabahar*-CgCOM1*i) and tomato-RNAi (Pusa Rohini-*CgCOM1i* and Pusa Early Dwarf-*CgCOM1i*) lines using fungal spore inoculum (1×10^5^spores ml^−1^), prepared from 3-days old cultures as described above. Empty vector transformed (EV) and wild type (WT) plants challenged with the fungal pathogen served as controls. Inoculation in the case of leaves was performed on abaxial surface. And in the case of fruits, surface of the fruit was first pricked with a toothpick and then inoculum was applied. Infected leaves and fruits were incubated at 25 °C inside a humidified plastic box, and allowed for the development of the disease and progression of lesion formation. For assessing the progression of disease symptoms, leaves and fruits were photographed at regular intervals (0-10 days post inoculation) and used for measuring infected area with Image-J analysis program (https://imagej.nih.gov/ij). Experiments were repeated three times with 5 biological samples each. Percentage of infected area in leaf and fruit was calculated as the infected leaf or fruit area/total leaf or fruit area X 100. The t-test was performed on calculated infected leaf or fruit areas (%) for determining significance (p≤0.0001).

Infected tissues of leaf and fruit samples from *Cg*-infected chilli- and tomato-*CgCOM1*i plants and the control (EV and WT) plants were collected at different time intervals (at 0, 7, 36, 120 hpi and 9 dpi) and used for determining *CgCOM1* expression levels. For these studies total RNA was isolated from the infected tissues, cDNA prepared and used for assessing *CgCOM1* transcript abundance by semi-quantitative RT-PCR and quantitative RT-qPCR using appropriate primer sets (Supplementary Table 1). Simultaneously, expression levels of internal control genes, *C. gloeosporioides β-TUBULIN, CaGAPDH* in the case of *C. annuum* (Chilli) and *SlACTIN* in the case of *S. lycopersicum* (tomato) were also estimated using gene specific primers (Supplementary Table 1) Expression levels of *CgCOM1* as determined in quantitative and semi-quantitative RT-PCR studies were normalized with the expression levels of internal control genes. In RT-qPCR investigations, experiments were performed with two biological and three technical replicates, and fold changes in gene expression were computed by 2^-∆∆Ct^ method (Livak and Schmittgen, 2001).

### Microscopy of infected leaves

Staining of infected tissues was done according to Koch *et al.* (2013). Infected leaves were collected at different time intervals post-inoculation (dpi), transferred to microfuge tubes (2 ml) having 1 ml of Trypan blue solution, and boiled for 3 min. Subsequently, the leaves were destained overnight in chloral hydrate solution, and observations were scored with Image analyzer (EVOS^®^ FL, Thermo Scientific, USA).

## Results

### *Identification of* CgCOM1 *gene, phylogenetic analysis and the development of RNAi construct*

In ascomycetous *Magnaporthe oryzae*, insertional inactivation as well as deletion of *COM1* (*MoCOM1*) gene produced defects in conidial development, appressoria formation, penetration and infectious growth resulting in significantly reduced virulence of this fungal pathogen in rice and barley plants (Bhadauria *et al.*, 2010; Yang *et al.*, 2010).

*MoCOM1* gene encodes for a helix-loop-helix type transcription regulator, which is unique to filamentous ascomycetes (Supplementary Fig. S2). CgCOM1 (accession no. L2FQN8) protein from *Colletotrichum gloeosporioides* Nara gc5 showed about 36% identity and 52% similarity at amino acid level with MoCOM1 (Q45VP6). Database searches revealed that homologues of MoCOM1 are widespread in ascomycetous fungi. Hence, phylogenetic analysis of COM1 was carried out to determine evolutionary relationship of CgCOM1 with the homologues derived from other *Colletotrichum spp.* and *M. oryzae* (Supplementary Fig. S3). From among the species analyzed, COM1 proteins from *C. gloeosporioides* (L2FQN8), *C. tofieldiae* (A0A161Y4N1), *C. incanum* (A0A167E1D2), *C. orbiculare* (N4V176), and *C. higginsianum* (A0A1B7YD68) belonging to the family Glomerellaceae showed maximum resemblance with highly conserved protein sequences (76-78% identity at amino acid level). Pair-wise alignment of COM1 proteins from *Colletotrichum* spp. exhibited closer resemblance with COM1 homologues of *Verticillium alfalfae and V. dahlia* (47-54% identity) belonging to the family Plectosphaerellaceae, than *M. oryzae* COM1 (36% identity). Protein sequence analysis (InterPro) revealed that COM1 protein of *C. gloeosporioides* has conserved winged helix DNA binding domain (218-290 aa) and shares structural similarities and common evolutionary origin with *M. oryzae* COM1 protein (Supplementary Figs. S3 and S4).

Taking a cue from the studies on the role played by *COM1* in developmental and infection processes of *M. oryzae* (see above), we hypothesized that *COM1* gene in *C. gloeosporioides* also likely plays a similar role in contributing to infective capability and virulence of this ascomycetes fungal pathogen as well. Hence, silencing of its expression may lead to disruption in conidial formation and infectious growth, thus restricting spread of anthracnose disease. So in current study we aimed to interrupt the development of *C. gloeosporioides* by Host-Induced Gene Silencing of *CgCOM1* to prevent infection, with onset and spread of anthracnose disease in chilli and tomato. For this purpose, *CgCOM1* gene sequence of *C. gloeosporioides* was utilized for the development of *CgCOM1* silencing vector. For generation of RNAi construct we have selected 168 bp fragment (1849 nt to 2016 nt) of *CgCOM1* gene (XM_007282173) (Supplementary Fig. S1) having minimal homology to plant genes to avoid off-target effects. The selected *CgCOM1* gene fragment was amplified using gene specific primers (Supplementary Table S1) and cloned into pBlueScript SK. For developing the RNAi cassette, the *CgCOM1* gene fragments were excised from pBlueScript SK and cloned into RNAi plant transformation vector pMVR-hp (Reddy and Rajam, 2015), downstream of the Cauliflower Mosaic Virus 35S promoter, in sense and antisense orientations on both sides of the chalcone synthase intron spacer (Supplementary Fig. S4).

### *Generation of* CgCOM1-*RNAi chilli and tomato transgenic lines*

Transgenic plants of chilli (*Capsicum annuum* varieties, Pusa Jwala – PJ and Pusa Sadabahar PSB) and tomato (*Solanum lycopersicum* varieties, Pusa Rohini – PR and Pusa Early Dwarf PED) were generated via *Agrobacterium*-mediated transformation using A. *tumefaciens* LBA4404 harbouring pMVRhp-*CgCOM1* construct. A total of 42 putative transgenic plants of PJ and 60 plants of PSB in chilli (Supplementary Fig. S5; Supplementary Table S2), and 27 plants of PR and 22 plants of PED in tomato were generated from independent explants (Supplementary Fig. S6; Supplementary Table S2). Among these plants, integration of *CgCOM1*-RNAi gene was confirmed in 34 plants of PJ and 30 of PSB in chilli, and 22 plants of PR and 17 of PED in tomato through PCR using *CaMV35S* promoter- and *CgCOM1* gene-specific primers (Supplementary Figs. S7 and S8; Supplementary Table S2).

### Transgene expression and siRNA production

Relative expression levels of *CgCOM1* transgene in leaves of 10 PCR positive transgenic lines each of PJ and PSB of chilli and 5 transgenic lines each of PR and PED of tomato were analyzed by RT-qPCR. The RT-qPCR analysis showed variations in *CgCOM1* expression levels in different transgenic lines (Fig. 1, Supplementary Tables S3 and S4).

**Fig. 1.**
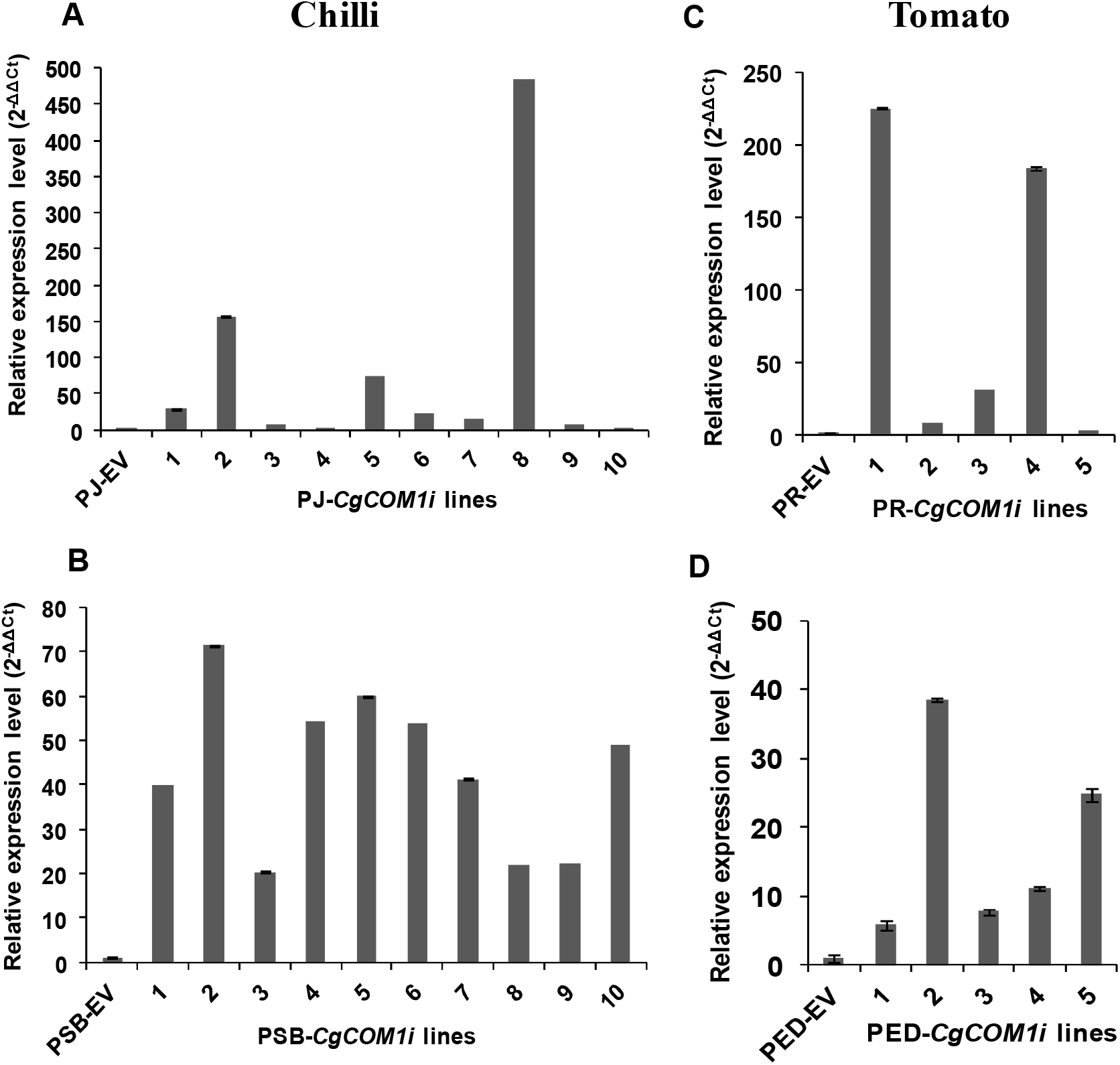
RT-qPCR analysis showing relative levels of *CgCOM1* gene transcripts in young apical leaves of RNAi transgenic chilli (A, B) and tomato (C, D) lines in comparison to respective vector control plants. Relative levels of target gene transcripts in the RNAi lines were normalized with *GAPDH* and *ACTIN* (internal controls) expression levels in chilli and tomato, respectively. Error bar represents mean ± SE with the three replicates. Relative expression of gene was computed by 2^-∆∆Ct^ method. PJ - Pusa Jwala, PSB - Pusa Sadabahar, PR - Pusa Rohini and PED - Pusa Early Dwarf; Vector control plants-PJ-EV, PSB-EV, PR-EV and PED-EV. Values on X-axis represent line numbers.

To verify the production of small interfering RNA (*CgCOM1-*siRNA) in *CgCOM1*-RNAi transgenic plants of chilli and tomato, stem-loop RT-PCR was carried out with cDNA of leaf tissue using siRNA specific forward and universal reverse primers (Supplementary Table S1). In *CgCOM1*-RNAi transgenic plants of both chilli and tomato, the stem-loop RT-PCR generated an amplicon of about 60 bp long, while no amplification was observed with control plants (Supplementary Table S3; Supplementary Fig. S9).

### Challenge assays on leaves and fruits of chilli and tomato RNAi lines

In order to assess resistance of transgenic plants against anthracnose disease, challenge assays were performed with *C*. *gloeosporioides* against detached leaves and fruits derived from the *CgCOM1*-RNAi chilli (PJ-*CgCOM1i* −1, −3, −9 and PSB-*CgCOM1i* −1, −4, −6) and tomato (PR-*CgCOM1i* −1, −2 and PED-*CgCOM1i* −1, −5) transgenic lines as described in materials and methods. Observations on spread of disease were scored at regular time intervals, and represented as percentage values of lesion areas compared to that in control plants (Figs. 2 and 3). Wild-type (WT) and empty vector (EV) transformed plants served as controls.

**Fig. 2.**
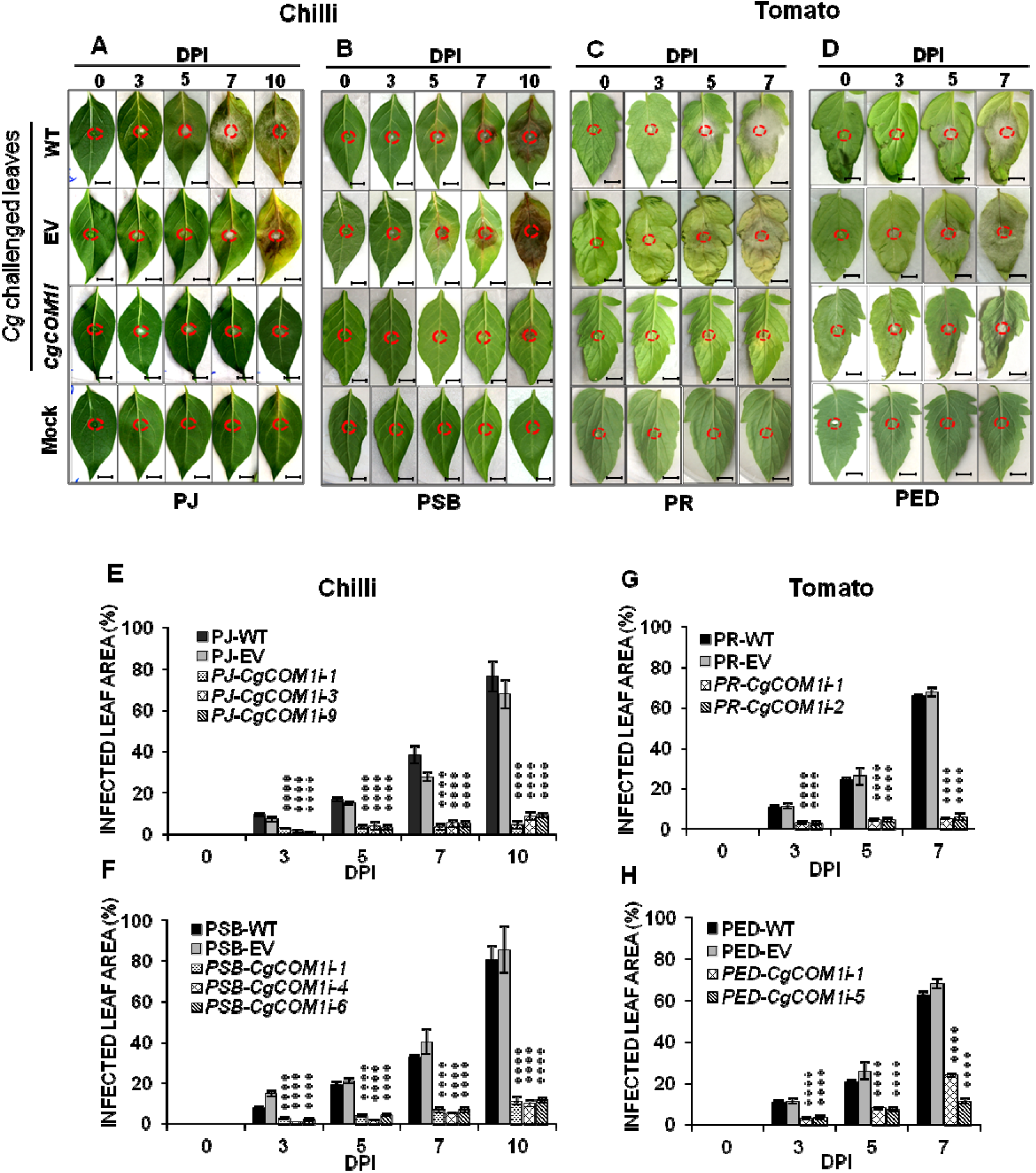
Infection symptoms and disease progression in *C. gloeosporioides-*challenged leaves of chilli (A and B) and tomato (C and D), and percent diseased / infected area in chilli (E and F) and tomato (G and H) RNAi lines relative to vector control and WT plants at 0, 3, 5, 7 and 10 DPI (Days Post Inoculation). Reduction in infection progression on *CgCOM1*-RNAi leaves in comparison to EV and WT leaves was highly significant (**** p ◻ 0.0001; Student’s t test). Red circles indicate sites of inoculation with *C. gloeosporioides* spores (challenged) or sterile water (mock). Bar scales in A-D represents 1 cm. Error bars in E-H represent mean values ± SE of three independent experiments. PJ - Pusa Jwala, PSB - Pusa Sadabahar, PR - Pusa Rohini and PED - Pusa Early Dwarf; Vector control plants-PJ-EV, PSB-EV, PR-EV and PED-EV.

**Fig. 3.**
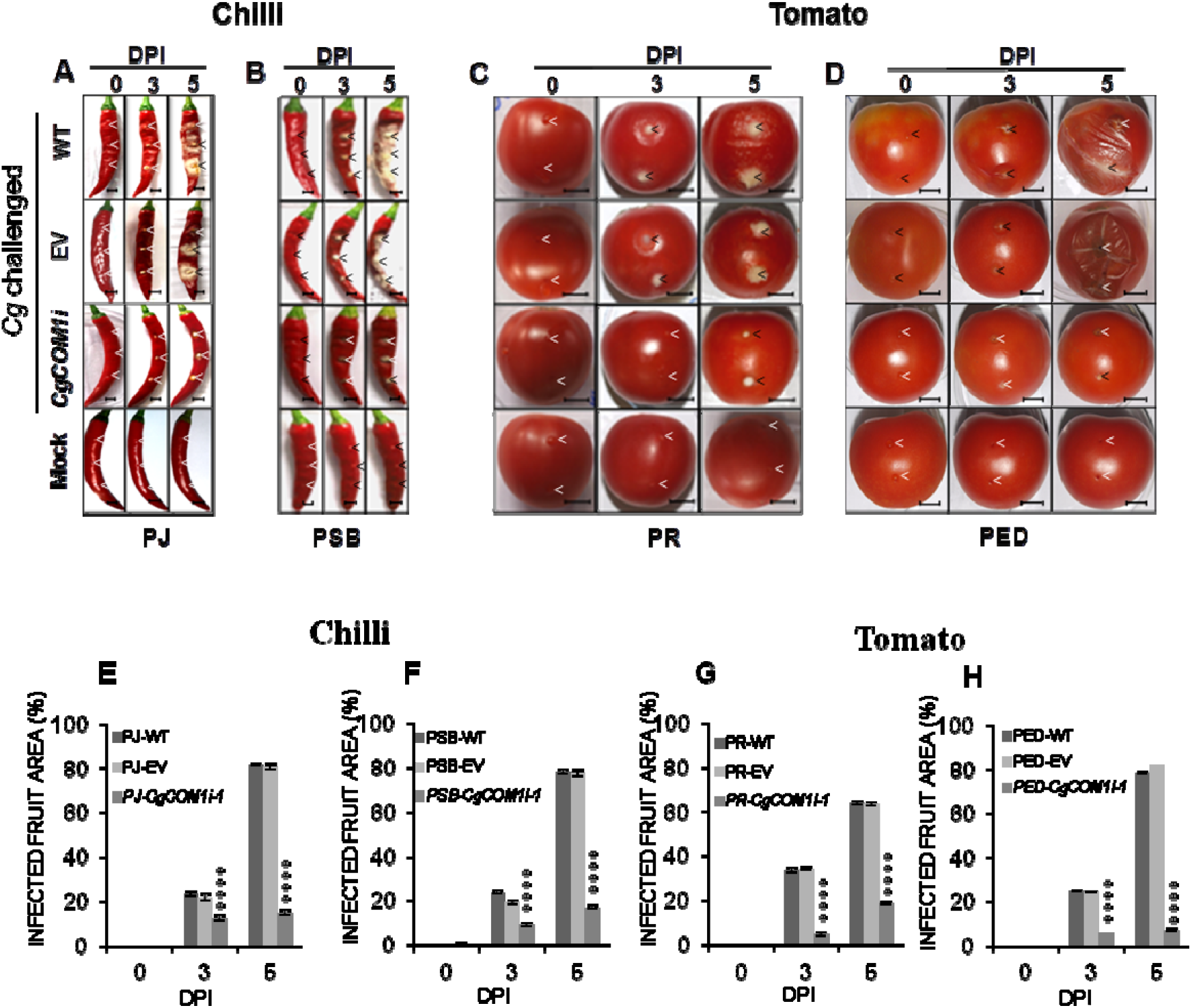
Infection symptoms and disease progression in *Cg-*infected fruits and quantification of infection area in chilli and tomato RNAi lines, EV and WT at 0, 3, and 5 DPI (Days Post Inoculation): Infected fruits and percent increase in infected area in PJ (Pusa Jwala, A and E), PSB (Pusa Sadabahar, B and F), PR (Pusa Rohini, C and G) and PED (Pusa Early Dwarf, D and H). Scale bar represents 0.5 cm (A, B) and 1.5 cm (C, D). In E-F, error bar represents mean value ± SE of three independent experiments. Reduction in infection symptoms on *CgCOM1i* expressing fruits compared to empty vector (EV) transformed and wild type (WT) control plants is highly significant (****P ◻ 0.0001; Student’s t test). Inoculum was introduced by pricking the fruit (arrowed). Mock: Inoculated with sterile water.

#### Assays on transgenic leaves

Chilli and tomato *CgCOM1*-RNAi lines challenged with the fungal pathogen showed extremely low levels of lesion development in leaves due to highly impeded growth of the fungus and spread of disease, as compared to wild type and vector control plants (Fig. 2). In wild type and vector control chilli and tomato leaves, water soaked spots with necrotic lesions, typical of anthracnose disease symptoms caused by *C. gloeosporioides*, appeared by 3 DPI. In contrast, by 3 DPI, the challenged leaves from *CgCOM1*-RNAi lines of chilli and tomato showed extremely low levels of disease symptoms (Fig. 2A–D). In chilli, percentages of infected areas on the leaves at 3 DPI was found to be 2.8%, 1.3% and 0.9% in the PJ-*CgCOM1i* lines 1, 3 and 9, respectively, as compared to 7.26% in vector control plants (Fig. 2E). Similarly, in the PSB-*CgCOM1*i transgenic lines 1, 4 and 6, lesion areas in leaves attained only a maximum of 2.6%, 1.3% and 2%, respectively, as compared to 14.7% in vector control plants at 3 DPI (Fig. 2F). By 10 DPI, disease symptoms in the *Cg*-challenged *CgCOM1*-RNAi lines of both the chilli varieties remained significantly low, while the vector control plants exhibited extensive spread of disease (Fig. 2E, F); average infected area on leaves of the transgenic lines PJ-*CgCOM1i* was 7.3% and on the PSB-*CgCOM1i* lines it was 11%. On the contrary, leaves from vector control plants of PJ and PSB exhibited extensive disease symptoms with infected zones reaching up to 68% and 85%, respectively, of total leaf area (Fig. 2E, F).

In tomato, at 3DPI, infected area on the leaves of the PR-*CgCOM1*i transgenic lines 1 and 2 averaged around 3%, whereas in the PED-*CgCOM1*i transgenic lines 1 and 5 it was about 3.4%, in comparison to 11.2% and 11.5% in the vector control plants PR-EV and PED-EV, respectively (Fig. 2G, H). Progression of disease was slower in the transgenic lines of PR-*CgCOM1i* (average diseased area reaching only up to 5.7% of total leaf area) and PED-*CgCOM1i* (average infected area was around 17.6%) at 7 DPI as compared to the vector control plants, in which spread of disease was found to be very rapid -- infected area reaching as high as 68% of leaf area in PR-EV and 69.3% in PED-EV in a similar time-frame (Fig. 2G, H). Whereas, the uninoculated leaves of RNAi lines, EV control and WT plants (mock inoculated) showed no disease symptoms. Analysis showed that infection area (lesion showing the spread of the disease) in leaves of *CgCOM1*-RNAi chilli and tomato lines was about 65-93% lesser than in the WT and EV plants at 10 DPI and 7 DPI respectively (Fig. 2).

#### Assays on transgenic fruits

Ripened fruits of chilli and tomato *CgCOM1*-RNAi transgenic lines challenged with fungal pathogen showed extremely low levels of disease symptoms in comparison to wild type and vector control plants (Fig.3). Water soaked necrotic lesions accompanied by dense fungal mycelial growth appeared in wild type and vector control chilli and tomato fruits by 3 DPI, while *CgCOM1*-RNAi lines of chilli and tomato showed diminished lesion formation in comparison to EV and WT fruits (Fig. 3). Percentage of infected area on the fruits of PJ-*CgCOM1*i-1 transgenic lines was 12.44% compared with 21.89 % on the vector control plants (Fig. 3E). In PSB-*CgCOM1*i-1 transgenic lines it was 9.14% in comparison to 19.29% in vector control plants (Fig. 3F) at 3 DPI. Infected area at 5 DPI in transgenic lines was 14.76% in PJ-*CgCOM1i-1* and 16.87% in PSB-*CgCOM1i-1* in comparison to EV fruits where infected area increased to 80.57% in PJ-EV and 77.66% in PSB-EV. Disease in RNAi transgenic fruits reduced up to 81.69% in PJ-*CgCOM1i-1* and 78.28% in PSB-*CgCOM1i-1* compared to EV challenged plants (Fig. 3E, F).

The percentage of infected area in tomato fruits of PR-*CgCOM1*i-1 transgenic lines was 4.66% as compared with 34.23 % on the vector control at 3 DPI, while at 5 DPI the infection area increased up to 18.8% in PR-*CgCOM1*i-1 as compared to 63.57% in vector control fruits (Fig. 3G). Whereas, the percentage of *Cg*-infected fruit area on PED-*CgCOM1*i-1 transgenic lines was 5.89% compared with 24.46% on the vector control at 3 DPI and, at 5 DPI the infected area was 7.07% in PED-*CgCOM1*i-1 and 82.02% in vector control fruits (Fig. 3H). Disease progression in wild type and vector control fruits was almost similar.

### COM1 *gene of* C. gloeosporioides *is strongly silenced when inoculated onto* CgCOM1*i transgenic plants*

To validate the inhibition of disease progression is due to silencing of the *CgCOM1* gene, semi-quantitative RT-PCR and RT-qPCR analyses of *CgCOM1* in *Cg*-inoculated leaves and fruits of chilli- and tomato-*CgCOM1i* lines were performed at different time intervals. RT-PCR and RT-qPCR analyses revealed that the *CgCOM1* transcript levels are significantly reduced with time in the *Cg*-challenged chilli- and tomato-*CgCOM1i* lines as compared to EV plants (Figs. 4, 5 and Supplementary Fig. S10). In chilli leaf tissue, RT-qPCR analysis showed that *CgCOM1* transcript levels in *Cg*-infected *CgCOM1*i lines were reduced by about 9.5 to 9.9 folds (depending on the transgenic line) in PJ-*CgCOM1*i (Fig. 4A) and 6.8 to 10.5 fold in PSB-*CgCOM1*i (Fig. 4C) as compared to *Cg*-infected vector control plants at 120 hpi. On the other hand, in the *Cg*-infected chilli-*CgCOM1*i fruits, the reduction in transcript levels ranged between 9.3 to 10.8 folds in PJ-*CgCOM1*i (Fig. 4B) and 6.5 to 8.2 folds in PSB-*CgCOM1*i (Fig. 4D) compared to vector control plants at 120 hpi. In tomato, RT-qPCR analysis revealed that by 120 hpi, *CgCOM1* transcript levels in *Cg*-infected tomato-*CgCOM1*i leaves were low, by about 7.7 folds in the case of PR-*CgCOM1i*-1 (Fig. 5A) and 3.8 fold in PED-*CgCOM1i*-1 (Fig. 5B) compared to the *Cg*-infected vector control plants. In *Cg*-infected tomato-*CgCOM1*i fruits, the transcript levels of *CgCOM1* were down-regulated by about 8.5 fold in the case of PR-*CgCOM1i*-1 (Fig. 5C) and 9.8 folds in the case of PED-*CgCOM1i*-1 (Fig. 5D) in comparison to vector control plants.

**Fig. 4.**
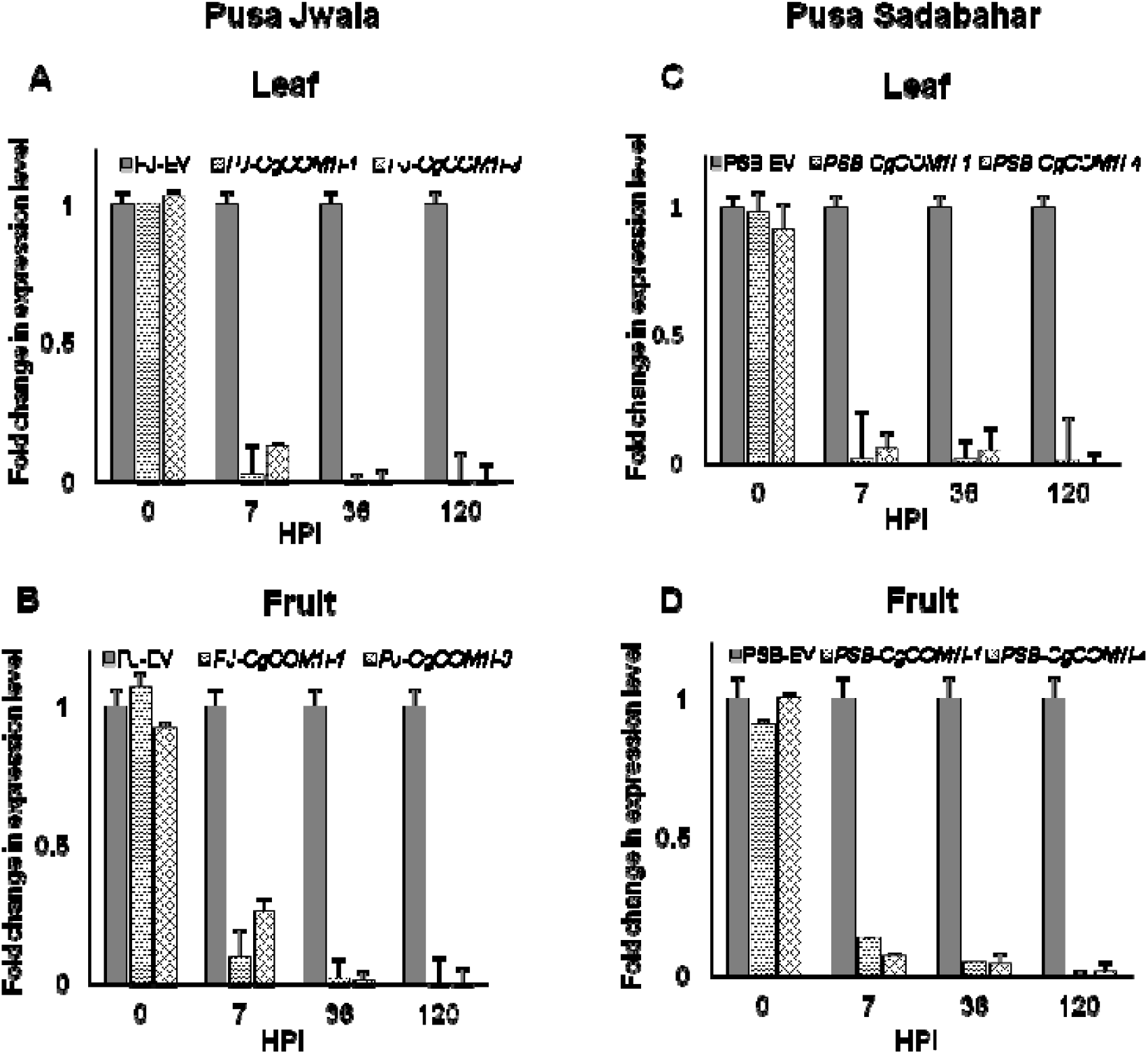
Host induced silencing of *COM1* gene in *C. gloeosporioides* in leaf and fruit tissues of chilli-RNAi plants. Transcript levels of the *CgCOM1* gene relative to the fungal housekeeping *TUBULIN* gene were determined by RT-qPCR analysis performed on RNA isolated at different time points (hours post inoculation) from the *Cg*-infected leaves and fruits of Chilli-RNAi lines and vector transformed plants (EV control). A-D. Represents expression levels in leaves (A, C) and fruits (B, D) derived from the *Cg*-infected PJ-*CgCOM1i*-1 and PJ-*CgCOM1i*-3 and PSB-*CgCOM1i*-1 and PSB-*CgCOM1i*-4 plants. Transcript levels were compared with the tissues collected at the same time points in both *Cg*-infected RNAi lines and vector control plants. Expression levels in *Cg*-infected RNAi plants were normalized against the challenged vector control plants and computed according to 2^-∆∆CT^ method. Error bars represent mean ± SE (standard error) values computed from two biological (three technical replicates each). (PJ - Pusa Jwala, PSB - Pusa Sadabahar; Vector control plants - PJ-EV and PSB-EV).

**Fig. 5.**
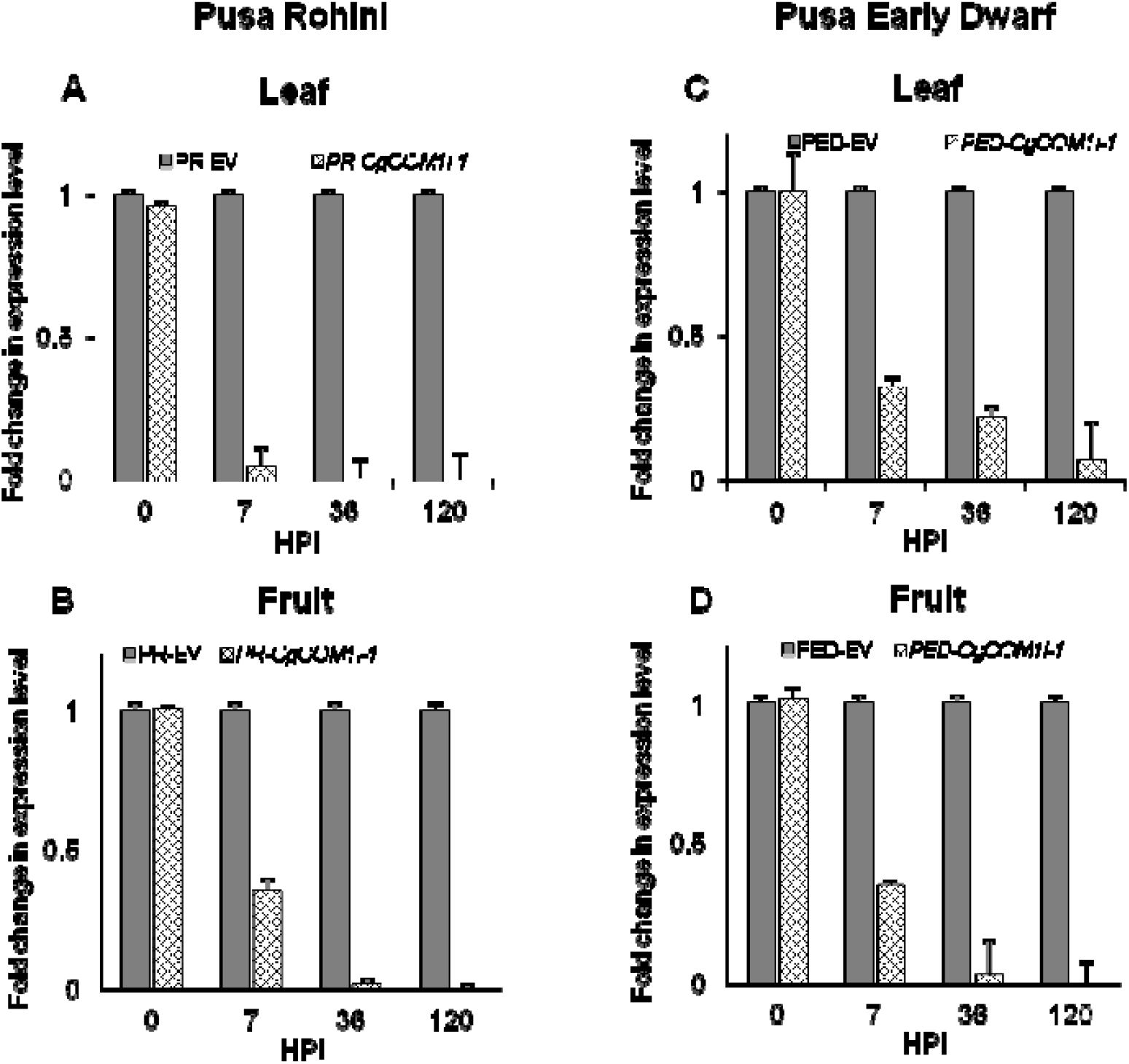
Host induced silencing of *COM1* gene in *C. gloeosporioides* upon infection of leaf and fruit tissues of tomato-RNAi plants. Quantification of expression levels of *CgCOM1* in the fungal pathogen infecting leaf and fruit tissues of tomato-RNAi lines was done by RT-qPCR analysis, similar to that performed with chilli-RNAi plants as described in Fig. 4. Expression levels in the leaves (A, C) and fruits (B, D) derived from the *Cg*-infected PR-*CgCOM1i*-1 and PED-*CgCOM1i*-1 plants. (PR - Pusa Rohini, PED - Pusa Early Dwarf; Vector control plants - PR-EV and PED-EV).

RT-qPCR analysis of *Cg*-challenged leaves and fruits of transgenic chilli and tomato RNAi lines demonstrated near complete silencing of the *COM1* gene in *C. gloeosporioides* by 120 hpi (5 dpi) compared with EV plants. Level of silencing of *CgCOM1* in the pathogen on *CgCOM1*-RNAi expressing leaves and fruits is in consonance with the degree of reduction in disease progression in plant tissues (Figs. 2–5).

### *Microscopic study of* C. gloeosporioides *infection process in* CgCOM1*i transgenic lines of chilli*

Our studies demonstrated a strong correlation between the reduced *CgCOM1* transcript levels as well as the lower incidence and manifestation of anthracnose disease in RNAi plants. In order to assess if the reduction in manifestation of anthracnose disease in RNAi plants is due to impairment of the fungal development caused by the plant derived *CgCOM1* siRNA, we performed microscopic evaluation of the fungal growth and development on *CgCOM1*i plant tissues. Under *in vitro* growth conditions on PDA (Potato Dextrose Agar) medium, germination of conidia of *C. gloeosporioides* commenced after 3 hpi, and by 18 hpi almost all spores germinated. Generally appressorial formation was observed by 36 hpi at a low frequency (Supplementary Fig. 11). Germination pattern of the fungal pathogen on the leaves of *Cg*-challenged vector control plants was similar to that found in *in vitro* conditions excepting that by 36 hpi hyphal growth was extensive with profuse differentiation/proliferation of appressoria (Fig. 6). In addition, on *Cg*-inoculated control leaves, nectrotrophic growth of *C. gloeosporioides* was initiated by 36 hpi. In contrast, in *Cg*-inoculated leaves of *CgCOM1*i plants, conidial germination at 36 hpi was found to be significantly lower with impeded germ tube growth and appressoria formation (Fig. 6). Repression of germ tube growth lead to curtailed hyphal development, and appressorial differentiation culminating into lobed or wrinkled appressoria. By 120 hpi, hyphal growth on the *Cg*-inoculated leaves of *CgCOM1*i plants was found to be highly impeded, with virtually no appressoria formation (Fig. 6). Hence, we deduce that the *CgCOM1* gene silencing in *Colletotrichum*, caused by the *CgCOM1*-siRNA produced by the host plant, is resulting in the development of non-functional infection-apparatus leading to the loss of pathogenicity of the fungus.

**Fig. 6.**
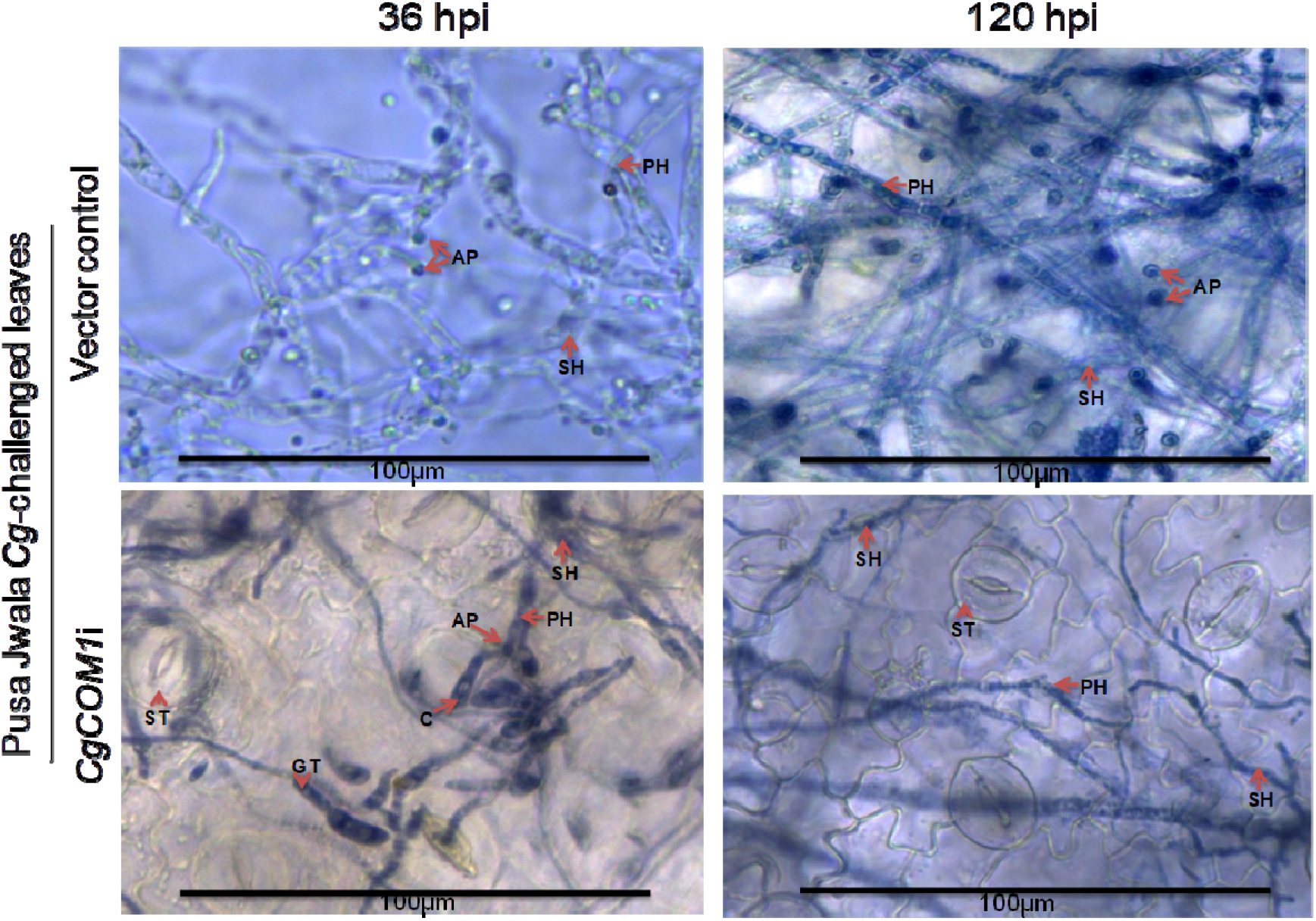
Photomicrographs showing mycelial growth and disease progression on abaxial surface of the *C. gloeosporioides*-challenged leaves of Pusa Jwala-*CgCOM1*i and vector control chilli plants at 36 and 120 hours post inoculation. Images were taken at (A) 10X and (B) 20X magnification using EVOS-FL imaging system. C - Conidia, AP - Appressorium, GT - Germ tube, PH - Primary hyphae, SH - Secondary hyphae and ST - Stomata. Sizes of bar scales are represented at the bottom of micrographs.

## Discussion

Since last ten years, HIGS strategy is widely employed to control plant diseases via suppression of pathogen infection by targeting and silencing a variety of pathogen genes to disable pathogen development/differentiation and pathogenicity (Bharti *et al.*, 2017; Chen *et al.*, 2016; Ghag *et al.*, 2014; Jahan *et al.*, 2015; Koch *et al.*, 2013; Nowara *et al.*, 2010; Song and Thomma, 2018; Thakare *et al.*, 2017). These studies showed that selection of appropriate target gene(s) is the most important requirement for developing an effective HIGS to incapacitate fungal pathogens. For example, Nowara *et al.* (2010) using susceptible barley (*Hordeum vulgaris*) and wheat (*Triticum aestivum*) cultivars demonstrated that severity of the powdery mildew fungus *Blumeria graminis* could be curtailed through HIGS by expressing dsRNA of the fungal *Avra 10* gene in plants. In 2013, Koch *et al.* employing HIGS strategy in Arabidopsis and barley targeted the fungal lanosterol C-14α-demethylase (*CYP51A, CYP51B* and *CYP51c*) genes to incapacitate the pathogen *Fusarium graminearum*, thus conferring the plants complete resistance against *Fusarium* head blight disease. Host-mediated silencing of two vital fungal genes (*Velvet* and *Fusarium transcription factor 1*) of *Fusarium oxysporum* was shown to confer banana plants resistance against *Fusarium* wilt disease (Ghag *et al.,* 2014). On lettuce (*Lactuca sativa*), growth of a biotrophic oomycete (*Bremia lactucae*) that causes downy mildew could effectively be suppressed by the host induced silencing of *Highly Abundant Message #34* (*HAM34*) and *Cellulose Synthase* (*CES1*) genes (Govindarajulu *et al.,* 2015). Cheng *et al.* (2015) successfully improved resistance of wheat against *Fusarium graminearum* pathogen that causes *Fusarium* head blight and *Fusarium* seedling blight by silencing of the fungal virulence gene *chitin synthase 3b* (*Chs3b*) through HIGS. Similarly, Bharti *et al.* (2017) demonstrated host induced silencing of *FOW2* and *chsV* genes in *Fusarium oxysporum* in transgenic tomato plants conferring enhanced resistance to vascular wilt disease. Recently, Song and Thomma (2018) through host-induced silencing of *Ave1, Sge1* and *NLP1* genes of *Verticillium dahliae* created tomato and Arabidopsis plants resistant to *Verticillium* wilt disease.

In *M. oryzae,* mutation in *COM1* gene significantly hindered conidial development and appressorial formation, thus disabling proper conidial germination and pathogenicity of the fungus in rice (Bhadauria *et al.*, 2010; Yang *et al.*, 2010). Based on the critical role played by *COM1* gene in promoting infection of *M. oryzae* in rice, for our studies we have selected *COM1* homologue as a potential target for host induced gene silencing in *C. gloeosporioides*. In order to neutralize the growth of the fungal pathogen *C. gloeosporioides* by HIGS, we developed chilli and tomato transgenic plants carrying hairpin *CgCOM1* RNA expression cassette and showed that the majority of infected *CgCOM1i* transgenic chilli and tomato lines are able to effectively inhibit the progression of anthracnose disease due to the trans-expression of *CgCOM1* siRNA in leaves and fruits; displaying up to 90% resistance against *C. gloeosporioides* (Figs. 2–5). Even though the *COM1* transcript abundance in *C. gloeosporioides*-infected plant tissues progressively diminished to near about undetectable levels by 120 hpi, in some transgenic lines, at the initial stages up to 36 hpi, the presence of intact *COM1* transcripts albeit at low levels (before they were being completely degraded due to the action of *COM1* siRNA (Figs. 4 and 5) perhaps was enough to aid marginal progression of infection up to necrotrophic stage (Fig. 6). Inhibition of infection through HIGS of *CgCOM1* demonstrates that like in *M. oryzae*, *COM1* in *C. gloeosporioides* plays a critical role in pathogenesis of this fungal species in host tissues. More than 80% of the tested *CgCOM1i* transgenic lines showed elevated levels of resistance against *C. gloeosporioides* infection with minimal disease progression compared to the vector control and wild type plants.

Generally, conidia of *C. gloeosporioides* are aseptate, but when the germination is triggered and cell cycle is activated, septa are developed prior to the emergence of germ tube from conidia (Supplementary Fig. S11). Similar observations were recorded by Mould *et al.* (1991), O’Connell *et al,* (1993) and Moraes *et al.* (2013). Normally germinated conidia, germ tubes and appressoria adhered to the surface of the inoculated leaves (Fig. 6). In our studies with chilli, we always observed that infection of the leaf tissues occurred through penetration of epidermal tissue by invading hyphae of *C. gloeosporioides*, but not via stomatal openings (Fig. 6). Similar type of infection was also observed in guava (Moraes *et al.,* 2013). However, in cowpea leaves, it was reported that *C. gloeosporioides* attempt to gain entry into leaf tissues through stomatal openings (Eloy *et al.*, 2015). Under *in vitro* culture conditions on PDA medium, *C. gloeosporioides* typically exhibited growth stages like germination of conidia after 3 hpi and appressoria formation after 36 hpi; but never displayed the development of primary and secondary hyphae under these conditions. In contrast, on the leaves of *Cg*-challenged control chilli plants, conidial germination was extensive and it was followed by profuse appressorial differentiation and extensive hyphal growth, and initiation of the development of secondary hyphae (necrotrophic stage) by 36 hpi. According to Alkan *et al.* (2015) on tomato fruits, *C. gloeosporioides* conidia germinate and differentiate into appressoria within 6-19 hpi, and later appressoria give rise to thin penetration pegs at 40-48 hpi and finally secondary hyphae by 2-3 days. Studies of O’Connell *et al.* (2012) and Moraes *et al.* (2013) on lifestyle of *C. gloeosporioides* on host plants such as maize and guava also revealed similar growth patterns, but with slight variations in timings of developmental / infection stages in biotrophic (conidia germination, formation of appressoria, penetration peg and primary hyphae) and necrotrophic (formation of secondary hyphae) phases. Over all, above studies indicate similarities in developmental phases of *C. gloeosporioides* on various host plants.

In our study, RT-PCR and RT-qPCR analyses indicated that the expression of *CgCOM1* gene and level of it’s transcript abundance significantly increased with the passing of time in fruits (Supplemental Fig. S12) and leaves of wild type or vector control plants infected with the *C. gloeosporioides*. On the other hand, in the fungal pathogen infected tissues of chilli- and tomato-*CgCOM1*i transgenic plants, *CgCOM1* transcript abundance decreased to almost undetectable levels by 120 hpi (Fig. 4 and 5), indicating nearly complete degradation of *CgCOM1* transcripts via HIGS in the transgenic lines.

Our studies demonstrated that HIGS is a powerful tool to control anthracnose disease caused by *C. gloeosporioides* in chilli and tomato.

## Supplementary data

**Table S1.** Primers used in the present study.

**Table S2.** Generation of transgenic chilli (*C. annuum*) and tomato (*S. lycopersicum*) lines carrying *CgCOM1i.*

**Table S3.** Total number of chilli and tomato RNAi transgenic lines used in various experiments.

**Table S4.** Relative expression levels of *CgCOM1* gene in RNAi transgenic plants of chilli and tomato.

**Figure S1.** Nucleotide sequence of *Colletotrichum gloeosporioides* (strain Nara gc5) *COM1* gene fragment used for the preparation of RNAi construct.

**Figure S2.** Multiple sequence alignment of homologues of COM1 transcription factor from various *Colletotrichum* species and *Magnaporthe oryzae* using ClustalW program.

**Figure S3.** Phylogenetic analysis of *Colletotrichum gloeosporioides* COM1 protein.

**Figure S4.** (A) Schematic representation of T-DNA in pMVR-hp vector showing RNAi construct of *CgCOM1* and (B) Confirmation of pMVR-hp-*CgCOM1* construct by restriction analysis.

**Figure S5.** Generation of *C. annuum* transgenic plants.

**Figure S6.** Generation of *S. lycopersicum* transgenic plants.

**Figure S7.** Confirmation of transgene integration in *C. annuum* plants.

**Figure S8.** Confirmation of transgene integration in *S. lycopersicum* plants.

**Figure S9.** Stem-loop RT-PCR analysis to detect the presence of siRNA specific to the *CgCOM1* gene of *C. gloeosporioides* in chilli.

**Figure S10.** RT-PCR analysis depicting expression levels of *CgCOM1* gene in *C. gloeosporioides*-challenged vector control and *CgCOM1*i chilli leaves.

**Figure S11.** Spore germination and development of infective germlings of *C. gloeosporioides* under *in vitro* conditions.

**Figure S12.** Expression analysis of *CgCOM1* during various stages of infection in *Cg*-infected and uninfected wild type chilli fruits by RT-qPCR.

## Acknowledgements

The research work was financed by the Department of Biotechnology (Grant No. BT/PR5399/AGR/36/722/2012), Government of India, to SD and MVR. BKM gratefully acknowledges Department of Science and Technology-INSPIRE (DST-INSPIRE-IF120813), Government of India, for providing fellowship.

